# Decoupling growth and production by removing the origin of replication from a bacterial chromosome

**DOI:** 10.1101/2021.12.07.471534

**Authors:** Marje Kasari, Villu Kasari, Mirjam Kärmas, Arvi Jõers

**Affiliations:** Institute of Technology, University of Tartu, Nooruse 1, 50104 Tartu, Estonia

## Abstract

Efficient production of biochemicals and proteins in cell factories frequently benefits from a two-stage bioprocess in which growth and production phases are decoupled. Here we describe a novel growth switch based on the permanent removal of the origin of replication (*oriC*) from the *Escherichia coli* chromosome. Without *oriC*, cells cannot initiate a new round of replication and they stop growing while their metabolism remains active. Our system relies on a serine recombinase from bacteriophage phiC31 whose expression is controlled by the temperature-sensitive cI857 repressor from phage lambda. Reporter protein expression in switched cells continues after cessation of growth, leading to protein levels up to five times higher compared to non-switching cells. Switching induces a unique physiological state that is different from both normal exponential and stationary phases. Switched cells remain in this state even when not growing, retain their protein synthesis capacity, and do not induce proteins associated with the stationary phase. Our switcher technology is potentially useful for a range of products and applicable in many bacterial species for decoupling growth and production.

## Introduction

Production of value-added chemicals in living cells holds promise as a way to reduce the need for oil and divert to more sustainable production methods. The list of chemicals that can be produced through metabolic engineering grows constantly and there are hundreds, if not thousands, of scientific papers describing the biosynthesis route for some new product. However, only a small fraction of these are actually produced commercially and many bioproduction processes suffer from low yield, titre and productivity ^1^.

Forcing a growing cell to produce any product (protein or small molecule) at significant quantities inevitably generates a resource allocation conflict. Biomass increase and product formation both use the same general resources (carbon and energy) and usually also compete for the same key metabolite(s). This leads to a growth–production trade-off: high production strains grow slowly and rapidly growing strains have low product yields ^2,3^.

A two-stage bioprocess has been used to bypass this trade-off ^4^. In the first stage, cells are grown at maximal rate without any significant product production. At the desired moment, the growth is turned off and production is induced so that most of the available resources can be used for product formation. This two-stage bioprocess has been successfully implemented in strains where the switch from growth to production is part of their natural regulation. Combining an aerobic growth stage with an anaerobic production stage in *Corynebacterium acetoacidophilum* resulted in a high concentration of succinate ^5^ and a pH-shift-induced production stage in *Klebsiella pneumoniae* led to the accumulation of 2-ketogluconic acid ^6^. Other bacteria, such as *Escherichia coli*, do not have such natural regulation and their growth must be controlled by other means. Nutrient limitation (other than the main carbon source) has been used to curb *E. coli* growth while maintaining product synthesis ^7,8,9^. In one study, limiting nitrogen also inhibited carbon metabolism, but constraining phosphorus, sulphate or magnesium allowed cells to keep their metabolism active ^7^.

In recent years several artificial switches have been built to stop growth while leaving cells metabolically active. Brockman and Prather blocked fructose-6-phosphate utilization in glycolysis and channelled its precursor, glucose-6-phosphate, towards synthesis of myo-inositol ^10^. As a result, the growth rate of the culture decreased sixfold and myo-inositol production increased more than twofold. Klamt and colleagues utilized the temperature-controlled expression of an essential TCA cycle gene to stop growth ^11^; the resulting two-stage bioprocess showed an increase in titre and volumetric productivity of itaconic acid production. These “metabolic valve” approaches suppress some cellular pathway necessary for cell growth and channel carbon flow to the product production pathway. Lynch and colleagues combined phosphate limitation with a metabolic valve approach to limit cell growth and adjust its metabolism respectively. This led to the increased production of both small molecules ^12^ and proteins ^13^.

Li and colleagues used CRISPR/dCas9 to suppress expression of several endogenous genes, which resulted in growth reduction and increased expression of GFP ^14,15^. Notably, the best targets were several genes in the nucleotide biosynthesis pathway.

Chromosomal DNA replication in bacteria starts from the well-defined origin of replication, *oriC*. In *E. coli*, DnaA protein initiates replication by occupying its multiple DNA binding sites inside the *oriC* region and drives DNA unwinding ^16,17^. In subsequent steps, additional proteins are recruited, but the replication initiation is essentially controlled by the *oriC*–DnaA interaction.

Here we describe a novel approach to decouple growth and production, which does not require direct modification of any metabolic pathway. During the switch from growth to production, we induce the removal of *oriC* from the chromosome with the help of a site-specific serine recombinase. This prevents further initiation of chromosomal DNA replication and eventually cells stop growing. Cellular metabolism, however, remains active and cells keep expressing proteins long after cell growth has stopped.

## Results

### Temperature-induced excision of *oriC* in a switcher strain

We reasoned that the removal of *oriC* from the genome will stop replication initiation and lead to cessation of growth. To achieve this, we redesigned the vicinity of *oriC* in the *E. coli* genome by adding serine recombinase recognition and cleavage sites (*attB* and *attP*) on either side of it (Figure 1a). We also included a GFP reporter gene downstream of the *attB* site in such a configuration that GFP would be expressed only after excision of *oriC*; this allowed us to monitor cells in which there was successful recombination between *attB* and *attP* (i.e. switched cells) (Figure S1a).

**Figure 1.**
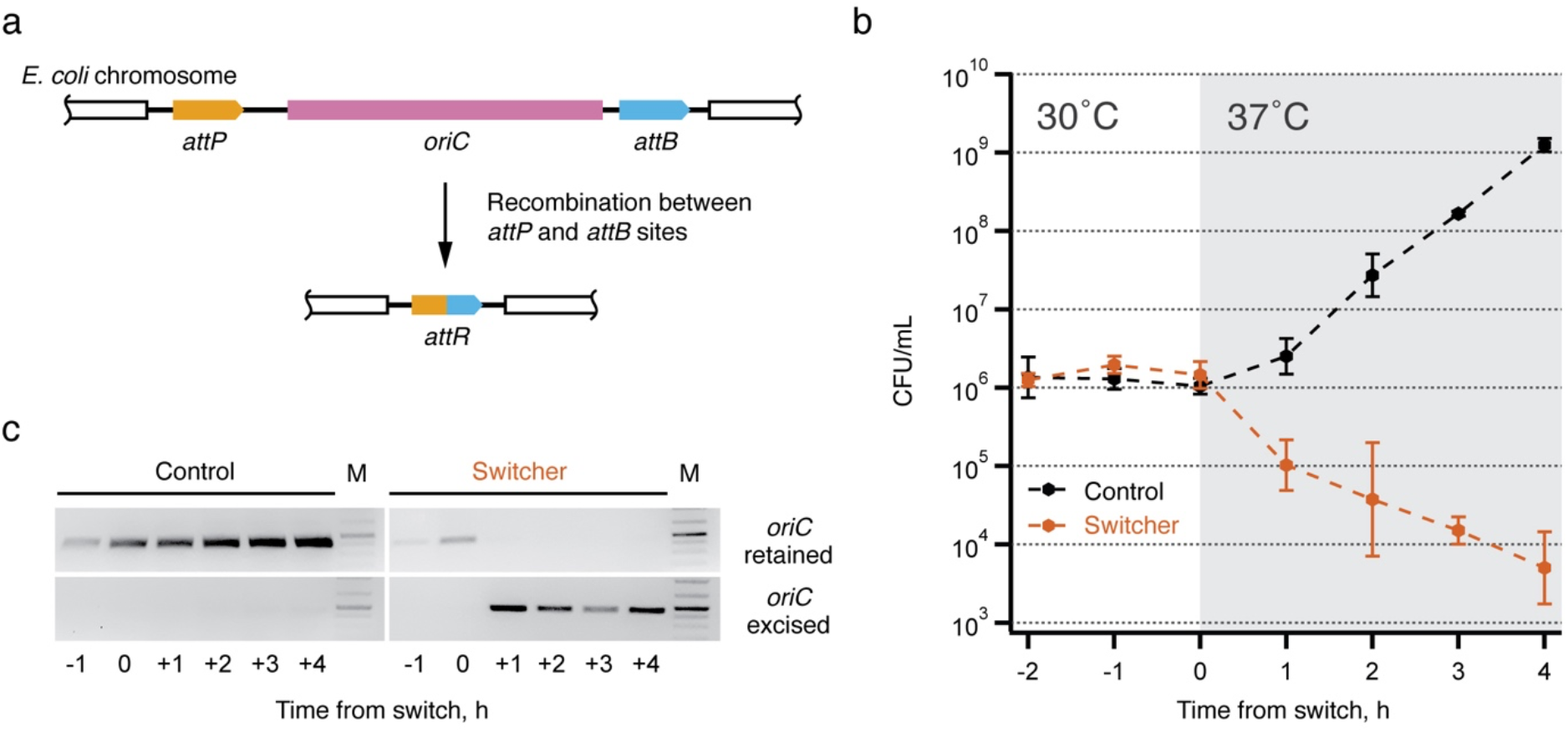
Expression of integrase leads to excision of *oriC* in the switcher cells. (**a**) Schematic setup of the switcher strain: serine recombinase recognition sites *attB* and *attP* are integrated either side of *oriC* in the bacterial chromosome in an orientation that leads to the excision of *oriC* upon recombination. (**b**) Control (black) and switcher (orange) cultures were pre-grown at 30°C (white plot area). At timepoint zero hours, the temperature was changed to 37°C (grey plot area). The number of colony-forming units was determined. The geometric mean of three independent experiments is plotted, error bars indicate standard deviation. (**c**) PCR was used to test switching on the chromosomal DNA using primers either unique to unrecombined (*oriC* retained) or recombined (*oriC* excised) DNA sequences and visualized using an agarose gel electrophoresis, M – DNA marker. Full uncropped gels are presented in Figure S2a.

The expression of a recombinase that is used to remove *oriC* from the genome must be tightly controlled to allow normal cell growth during uninduced conditions. We used the lambda phage cI857 transcriptional repressor ^18^ and its target promoter to control the expression of serine recombinase from bacteriophage phiC31 (phiC31 integrase) ^19^. cI857 is a temperature-sensitive mutant that represses its target promoter at 30°C, but the repression is relieved at 37°C. We generated an E. coli switcher strain in which initiation of the chromosomal replication can be eliminated. In its chromosome *oriC* sequence is flanked by phiC31-integrase-specific *attP* and *attB* sites and it carries a medium-copy expression plasmid pAJ35 encoding an inducible phiC31 integrase (Figure S1b, File S1). As a control strain, we used the same bacterial strain, but carrying plasmid pAJ27, which expresses only 20 N-terminal amino acids of phiC31 integrase and cannot catalyse recombination (Figure S1b, File S2).

To test the removal of *oriC*, we first measured the number of colony-forming units (CFUs) of switcher and control cultures grown at 30°C and 37°C. Non-growing bacterial cells cannot form visible colonies on a solid medium, and the lack of colonies is therefore an indication of the functional switch. The strains were precultured overnight at 30°C, diluted in the fresh medium and incubated at 30°C for 2 hours before shifting the temperature to 37°C. After the initial lag phase, the number of control culture CFUs started to increase as expected, whereas the switcher CFU count began to decrease (Figure 1b). Four hours after the temperature shift, the switcher CFU count is more than two orders of magnitude lower than at the time of the temperature shift, indicating the effective excision of *oriC*.

We verified the DNA rearrangement in switched cells by PCR amplification specific to either a pre-switch or post-switch DNA configuration (Figure 1c, Figure S2a). In the switcher culture, the pre-switch configuration is detected before the temperature shift, while a post-switch-specific PCR product appears after the culture has been shifted to 37°C. In control cells, only the product specific to the pre-switch configuration is present. We also verified the sequence of the *attR* site in the genome, which is formed during switching after recombination between *attB* and *attP* (Figure S2b).

### Switching enables the selection of final cell density

After removal of *oriC* from the genome, cells cannot initiate a new round of replication, and cell density should stabilize at a submaximal level. To test this hypothesis, we grew the switcher and control strains on a 96-well plate and followed the culture density before and after shifting the incubation temperature from 30°C to 37°C (Figure 2a). A few hours after the switch, the growth of the switcher strain attenuates, and its cell density eventually stabilizes at values different from a control strain. The plateau level reached depends on the cell density at the time of switch: the bigger the dilution from the overnight culture, the lower the plateau. Notably, the final density of a less diluted switcher culture (1600x) can even exceed the density of the stationary-phase control strain. Measured OD values of the control culture decrease through the stationary phase, likely because of cell shrinkage, whereas the decrease in switcher cultures is less prominent.

**Figure 2.**
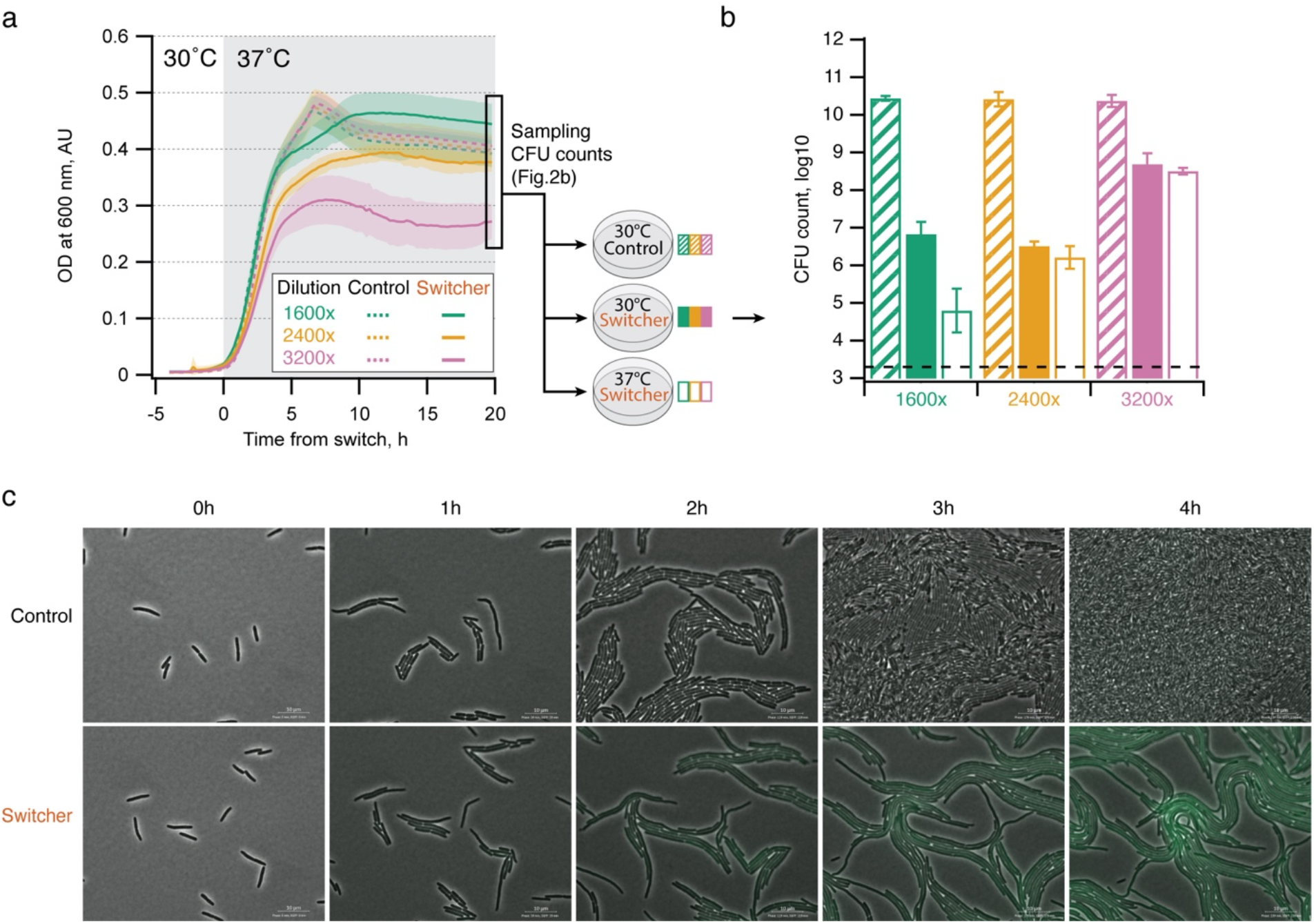
Switching enables the selection of final cell density. (**a**) The control strain (dashed lines) and switcher strain (solid lines) were inoculated from the overnight culture in various dilutions into fresh medium in a 96-well plate. At timepoint zero hours the temperature was changed from 30°C to 37°C to induce the switching. At the end of the experiment, the cultures were diluted and spotted to LB agar plates. (**b**) The number of colony-forming units was determined after incubating the plates for 20 hours at 30°C (control and switcher, striped or filled bars, respectively) or 37°C (switcher, empty bars). The dashed line indicates the detection limit. (**c**) Growth of control and switcher cells was monitored at 37°C for six hours using time-lapse microscopy. Hourly snapshots of cultures up to the 4 h timepoint are presented here. Full-length time-lapse videos are presented in Files S3 and S4. The mean of six biological replicates from two independent experiments is plotted on panel **a**; shading indicates standard deviation. The mean of three biological replicates is plotted on panel **b**; error bars indicate standard deviation.

To quantify the extent of switching we measured the number of CFUs in cultures after 20 h incubation at 37°C (sampled from the end of the experiment described in Figure 2a). Samples were plated on LB agar medium and incubated overnight at 30°C or 37°C as indicated on the figure. The switcher cells that have been incubated for 20 h at 37°C on a 96-well plate and still form colonies on an agar plate at 30°C have either not switched yet or have permanently lost the ability to switch (escaper mutants). On agar plates incubated at 37°C only escaper mutants can form colonies, meaning the removal of *oriC* is defective in these cells. Despite a similar culture biomass observed at the end of the experiment in Figure 2a (less than 2x difference), most cells in the switcher culture have switched, which can be seen because the number of CFUs on the 30°C plate has decreased by up to four orders of magnitude compared to CFUs of the control on the 30°C plate (Figure 2b). A fraction of escaper mutants seems to increase in a more diluted culture; in a 3200x dilution they constitute approximately 1% of all the cells in switching-favoured conditions.

Because the serine recombinase reaction is effectively irreversible, switched cells cannot return to growth even if the temperature is lowered back to 30°C degrees (as was tested in CFU measurements). This is different from growth regulation systems depending purely on gene activation and/or repression, because these cells can grow again after regulatory signalling is removed. Irreversible elimination of cell multiplication might be a useful feature for the development of living medicines, where bacterial cell growth must be kept under tight control ^20^.

To analyse the morphology of switching cells we used time-lapse microscopy. Cells were pre-grown at 30°C in LB medium and transferred onto an LB agarose pad under a microscope at 37°C. Both control and switcher cells keep growing for the first few hours (Figure 2c). Switcher cells start to express GFP, indicating switching has taken place. The switchers also become elongated and form filaments at the later timepoints, as seen in time-lapse videos (Files S3 and S4). This indicates that cell division is inhibited in switched cells.

Similar cell elongation have been shown by Nielsen and co-workers who used CRISPR-interference (CRISPR/dCAS9) mediated blocking of DnaA binding sites in *oriC* as a growth switch to generate a two-stage production system ^14^. This switch slowed down the growth but did not block it completely and resulted in a modest increase in protein production. Their approach was more efficient by targeting the genes *pyrF* ^14^, *pyrG* and *cmk* ^21^, leading to stabilization of culture density at a sub-maximal level. Notably, blocking of *pyrF* also led to elongated cells similar to those in our experiments (Figure 2c), indicating that this is a common reaction to the disruption of DNA metabolism.

Several examples of two-stage bioproduction techniques use metabolic valves, where the flow of metabolites is diverted from growth to production during the switch ^10,11,22^. While effective, these solutions are rather product-specific, and for almost every new type of product, a new growth switch must be built. Our switching approach blocks only DNA replication initiation and does not directly target other metabolic pathways, making it applicable for many products.

### Switching enhances protein expression

Stopping cell growth before all nutrients are depleted from a growth medium should leave more resources for protein production. We tested this assumption by inducing expression of a red fluorescent protein (mRFP1) at the time of the switch and by following mRFP1 fluorescence intensity (Figure 3a). Initially, mRFP1 accumulates in both control and switched cultures, but the accumulation stops in the control culture when it enters the stationary phase (OD curve plateaus). In contrast, switched cells continue to synthesise mRFP1 long after they have reached their maximum culture density. By the end of the experiment, the level of mRFP1 intensity is almost three times higher in the switched culture compared to the control, even when the cell density in the switcher remains lower.

**Figure 3.**
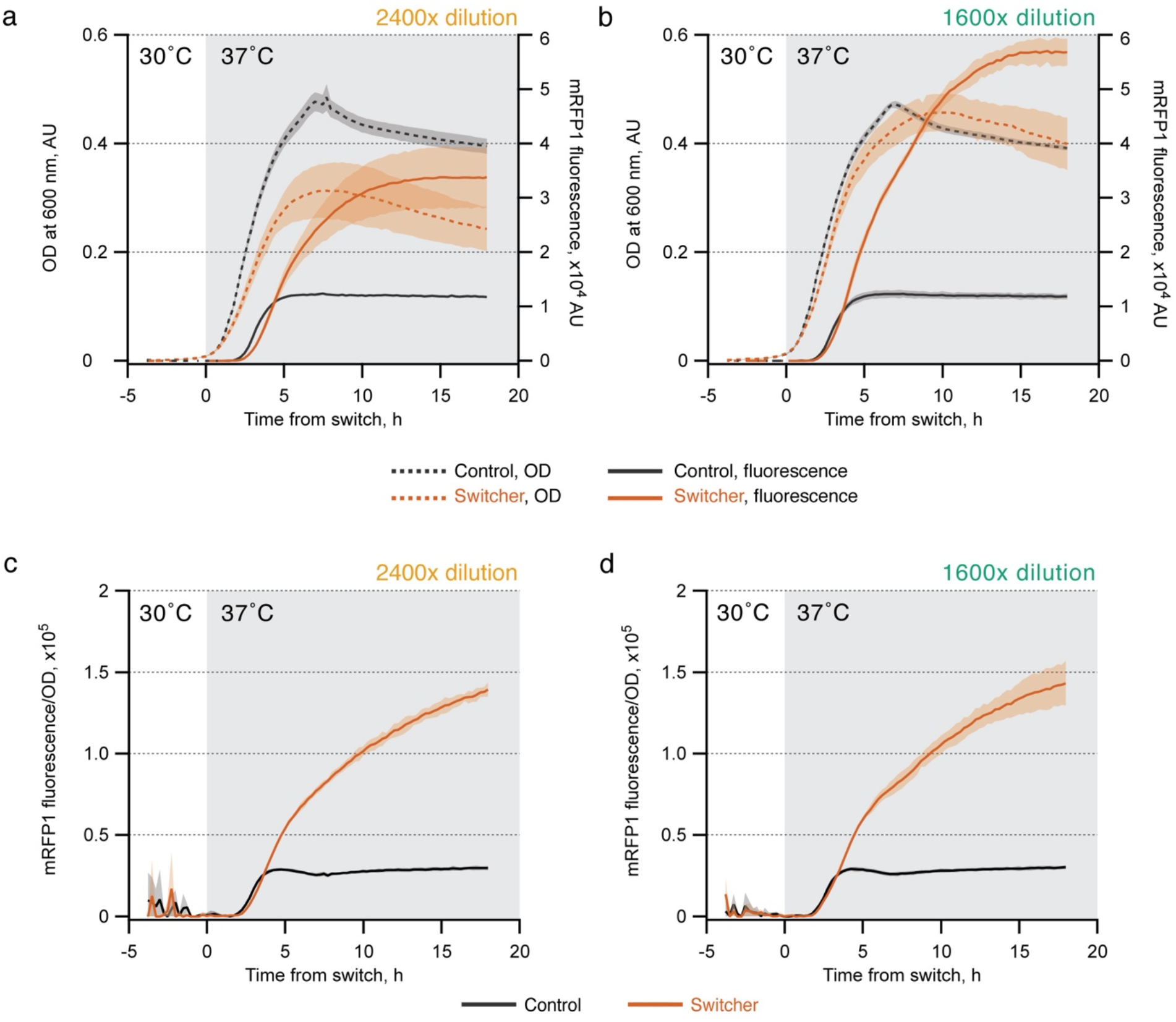
The assessment of protein production capability of the switcher and the control strains in LB. Overnight cultures were diluted 2400x or 1600x as indicated in the figure panels. Control (black) and switcher (orange) cultures were pre-grown at 30°C on a 96-well plate. At timepoint zero hours the temperature was changed to 37°C to induce switching. The production of fluorescent mRFP1 protein was induced by adding homoserine lactone (HSL) at timepoint zero. (**a, b**) The changes in OD at 600 nm (dashed lines, left axis) and increase in fluorescence (excitation: 584 nm, emission: 607 nm) of mRFP1 protein (solid lines, right axis) were monitored. (**c, d**) Fluorescence-over-optical-density ratio of control (black) and switcher (orange) culture was calculated based on values in panels **a** and **b**, respectively, and plotted. The mean of three biological replicates is plotted; shading indicates standard deviation.

The difference in protein expression is even more evident when the switching is timed so that the switched culture stabilizes at the same density as the control culture (Figure 3b). At the end of the experiment, the mRFP1 level in the culture with an initial 1600x dilution is five times higher than in the control. mRFP1 intensities normalized to OD are clearly higher in the switcher than the control and very similar in both dilutions (Figure 3c, d), indicating a more favourable protein-to-biomass ratio. To exclude a possible bias in one specific expression system, we tested protein synthesis capacity using a different reporter protein, promoter system, growth medium and plasmid backbone, and confirmed switcher-enhanced production of the protein of interest (Figure S3).

Changes in cell morphology, including filamentation, have been associated with increased accumulation of a desired product ^23^. To test if the elevated mRFP1 expression is dependent on cell elongation, we induced filamentation in the mRFP1-expressing cells by inducing SulA overexpression. SulA is a well-known FtsZ inhibitor and its expression in *E. coli* leads to cell filamentation ^24^. SulA-overexpressing cells become filamentous, but do not produce more mRFP1 (Figure S4), suggesting that reasons other than cell elongation are behind the increased protein expression in our switcher cells.

Lynch and colleagues used phosphorus limitation to constrain growth and metabolic valves to reconfigure metabolism. This separates growth limitation from metabolic rearrangement and allows use of the same growth-curbing mechanism for different types of products ^25,13^. Removal of *oriC* in our switcher is also a general way to stop growth and should be combined with appropriate changes in gene expression to achieve better production of the desired product. Both our switcher (Figure 3) and a phosphate-limitation-based system ^13^ allow protein accumulation after cell growth has stopped or significantly slowed down. This is in contrast to conventional protein expression where growth and production are coupled, and cells stop protein synthesis as soon as they reach stationary phase ^13^.

Coupled growth and production is problematic also for another reason. High-level production is a burden to cellular metabolism due to high resource usage and possible toxicity. This generates a selection pressure to accumulate cheater mutants – non-producer cells that have a growth advantage. Accumulation of cheaters is a well-recognised problem in bioproduction ^26^ and special effort in coupling production to the expression of an essential gene is needed to counteract the problem ^27,28^. When production induction is concurrent to, or initiated by, the growth switch, high-level production only occurs in switched cells, which will not grow anyway. Although some cells escape the switch and can grow at 37°C (Figure 2b), the frequency of mutants should be independent of burden, because growing cells do not express the product yet.

### Switched cells retain a protein synthesis capability long into the non-growing phase

Continuous accumulation of an expressed reporter protein despite unchanging cell density (Figure 3) prompted the question regarding how long switched cells can retain the ability to induce protein expression. To find this out, control and switcher cultures were grown on a 96-well plate and the expression of mRFP1 reporter was induced at different timepoints after the switch (Figure 4). Already three hours after the switch, induction in the control is very weak but almost unaffected in the switcher (Figure 4b). At 6 h, the control culture fails to induce any detectable mRFP1 expression, while in the switcher the mRFP1 induction is still strong. The addition of HSL, even after 20 or 30 h, results in mRFP1 induction, although at lower levels (Figure 4c). There is also a noticeable increase in mRFP1 signal in the uninduced switcher culture, which is probably caused by the partial derepression of the mRFP1 promoter. These results suggest that despite similar cell densities, the switched cells are in a different physiological state that is favourable for protein expression.

**Figure 4.**
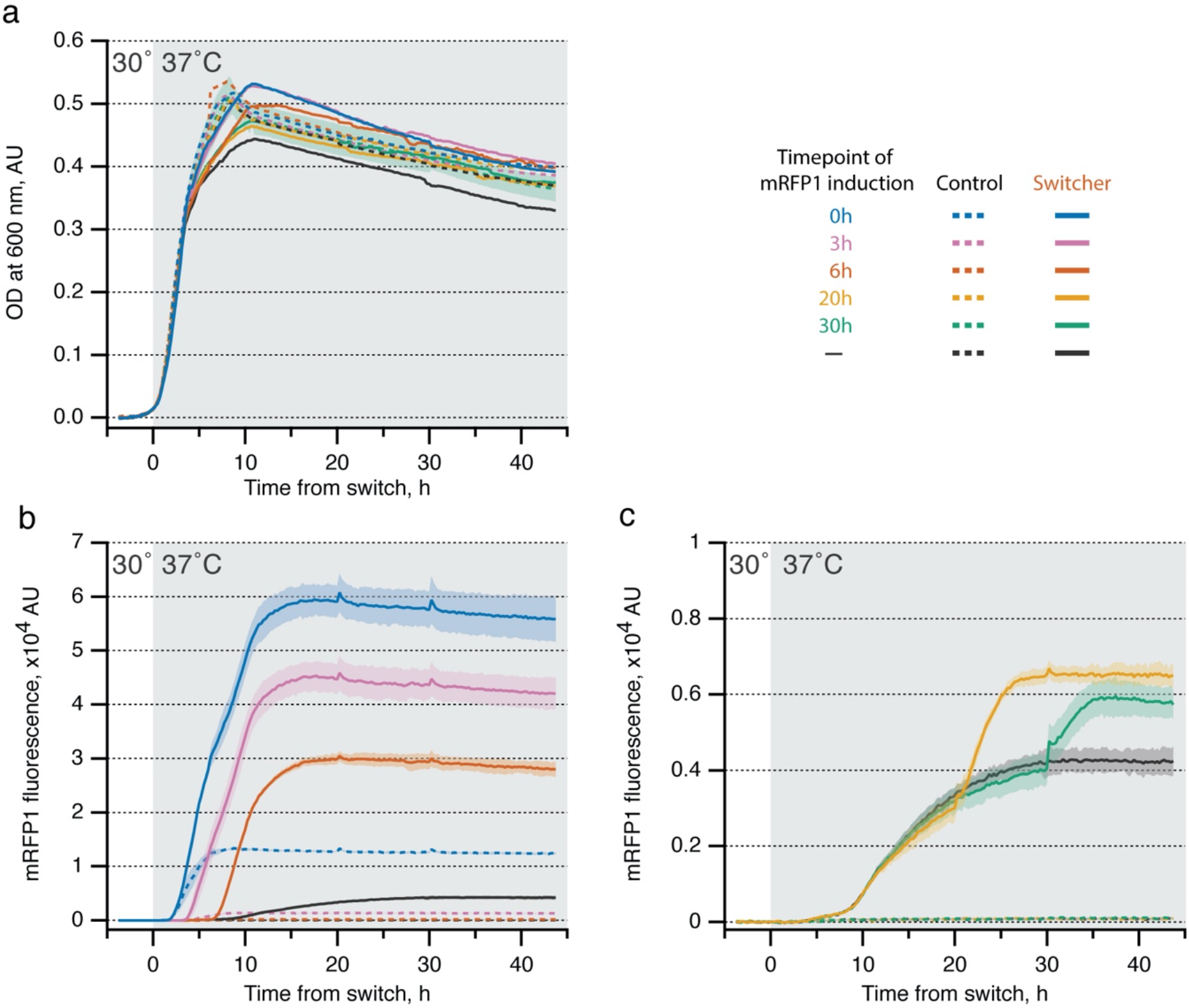
The protein production capability of the switcher remains active over a long period of time in LB. Control (dashed lines) and the switcher (solid lines) overnight cultures were diluted 1600x and pre-grown at 30°C on a 96-well plate. At timepoint zero hours the temperature was changed to 37°C to induce switching. The production of fluorescent mRFP1 reporter protein was induced by adding HSL at indicated timepoints. Changes in optical density (**a**) and fluorescence (**b, c**) (excitation: 584 nm, emission: 607 nm) were monitored. The mean of three biological replicates is plotted; shading indicates standard deviation.

### Switched cells enter into a state that is distinct from normal growing or stationary phase cells

Our results above indicate that non-growing switched cells are in a different physiological state compared to non-growing control cells (stationary phase cells). To characterize this state, we analysed and compared the proteome of switched and control cells. We designed the experiment so that the cell density in the switcher culture would either reach a similar level to the stationary phase of the control after the switch or stay at a significantly lower level. We collected samples of switched and control cells from growing, early-plateau, and late-plateau stages according to the growth curves of each culture, and additionally from a switched culture at low plateau (Figure 5a).

**Figure 5.**
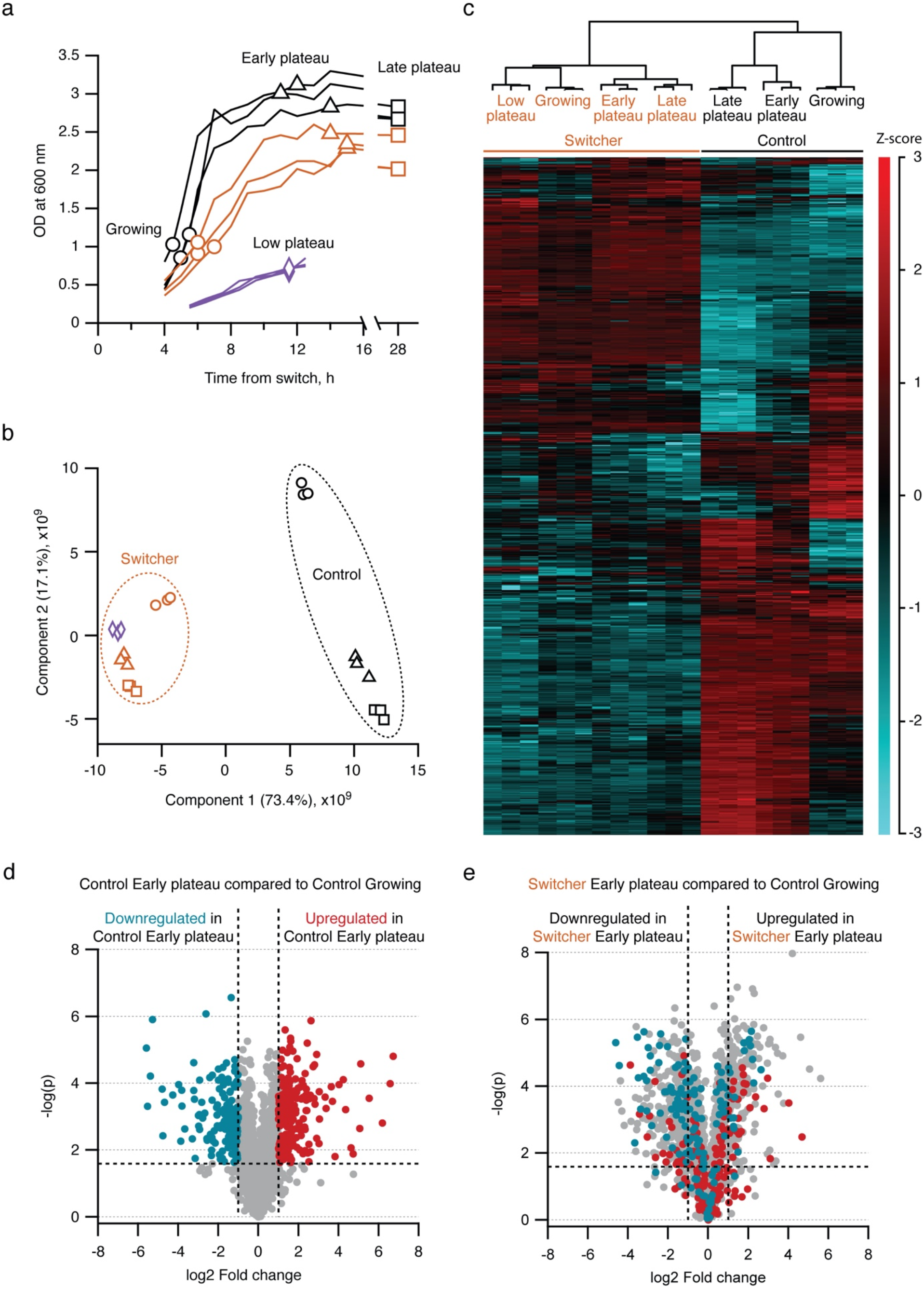
Comparison of proteomes of control and switcher cells. (**a**) Growth curves of control (black) and switcher (orange and purple) cultures. Two different initial dilutions from overnight culture were used to inoculate switcher cultures to obtain final cellular densities after the switch that were either similar to the control (orange) or approximately at OD ≈ 1 (purple). Sampling points for mass-spectrometric analysis are indicated with empty symbols. (**b**) Principal component analysis of switcher and control samples. (**c**) Hierarchical clustering heatmap of differentially expressed proteins. Proteins shown have statistically significant expression levels (multiple-sample ANOVA test false discovery rate (FDR) < 1%). Log2 transformed LFQ intensities were Z-scored across rows and hierarchical clustering using Euclidean distances was performed with the result presented as a heatmap. (**d**) Comparison of protein expression levels between control growing and control early-plateau samples. Red dots represent proteins upregulated in early-plateau cells, blue dots represent the downregulated ones (fold change > 2, FDR < 1%). (**e**) Comparison of protein expression levels between control growing and switcher early-plateau samples. Protein colour coding is the same as in panel **d**. The presented proteomic data was analysed using Perseus software and source data is available in File S10.

To create a low-dimensional representation of the variability between switcher and control samples, we performed a principal component analysis (PCA) using the whole proteome of each sample (Figure 5b). Proteome profiles of switched cells are different from both growing and stationary-phase (plateaued) control cells. All samples from switched cultures cluster together; growing and non-growing switched cells are notably more similar to each other than the respective samples of control cultures. Stable proteomic profiles support the hypothesis that switching leads to a distinct cellular state and that the switched cells stay in this state for at least 28 hours.

To analyse the differences between switcher and control cultures at the resolution of individual proteins, we visualized all statistically significant differences in protein abundance as a heatmap (Figure 5c). Statistical significance of changes in abundance of proteins between sample groups was determined by a multiple-sample ANOVA test, and hierarchical clustering of the significant proteins was performed on log2 transformed intensities after Z-score normalization. A hierarchical clustering dendrogram supports a similar sample group formation as in PCA and corroborates distinct differences between control and switcher cells. All samples from switched cultures cluster together and their protein expression pattern is clearly different from control samples.

We also analysed to what extent the non-growing switched cells express proteins that are associated with the stationary phase. We identified and colour-coded proteins in the control strain that are up- or downregulated in the early plateau (early stationary phase) and compared this to growing cells (exponential phase) (Figure 5d). Mapping these proteins on a plot comparing the switcher early plateau to growing control cells reveals that the stationary-phase-specific up- or downregulated proteins do not follow the same pattern in switched cells, but instead are distributed quite randomly (Figure 5e). We got similar results when comparing late-plateau samples of switcher to growing control cells (Figure S5a, b). To exclude that the distribution of up- and downregulated proteins in switcher samples is a result of random sample point variability we compared early- and late-plateau samples of the control culture (early and late stationary phase). Most of the proteins that are upregulated in the late stationary phase are also upregulated in the early stationary phase (Figure S5c).

Switched cells remain metabolically active even if a culture reaches the cell density normally characteristic of the stationary phase (Figures 3, 4). Corroborated by proteomics, switched cells are clearly different from stationary-phase cells (Figure 5). This suggests that the removal of *oriC* induces a state in which cells do not respond to signals that initiate the stationary phase in normal cells. Ribosome hibernation factors Sra, Rmf, Hpf and YqjD, which are responsible for inactivating ribosomes in the stationary phase ^29^, are not elevated in switched cells and display expression levels similar to growing cells (File S5). Our results also indicate that the lack of protein synthesis in the stationary phase is due to the physiological response and not because of a simple lack of carbon source or energy – the high-density switched culture kept synthesizing proteins, while a stationary-phase culture did not. Our switcher strain could, among other applications, be a useful tool to study the onset of the stationary phase and the role of chromosomal replication in it.

We identified proteins that are detected only in switcher samples and in none of the control samples, and those that are upregulated in all of the switched samples compared to each control sample (File S5). We mapped these proteins onto metabolic pathways using the toolset at EcoCyc.org and identified pathways that are upregulated in switched cells (p-value cut-off 0.05) (File S5).

Ribonucleoside reductases are responsible for dNTP synthesis, and their expression is elevated after switching (NrdA, NrdB, NrdD, File S5). DNA replication cannot be initiated in switched cells, and cells may perceive this as a lack of dNTPs. Transcription from the main ribonucleotide reductase operon *nrdAB* is stimulated by DnaA-ADP and inhibited by DnaA-ATP ^30^. Normally, DnaA-ATP level is high before DNA replication initiation and it is converted into DnaA-ADP soon after initiation. This cycle is absent in switched cells and a permanently low DnaA-ATP-over-DnaA-ADP ratio might be behind the elevated expression of Nrd proteins.

Enzymes participating in fatty acid synthesis pathways are also upregulated in switched cells. These include FabG, FabI, FabF, FabA, FabH and FabD, which are involved in several synthesis pathways. Their expression is controlled by FadR and FabR transcriptional regulators whose activity is regulated by unsaturated fatty acids ^31^. Notably, fatty acid synthesis can determine cell size in *E. coli* ^32^, so upregulation of fatty acid synthesis enzymes could contribute to the elongated cell phenotype.

Here we describe a novel way to decouple growth and production by removing *oriC* from the *E. coli* genome. We show how cells can remain metabolically active and keep synthesising proteins long after growth has stopped. For the purposes of bioproduction, our growth decoupling system could be combined with other measures that increase product synthesis, such as the up- and downregulation of specific metabolic pathways, substrates, or cofactor availabilities. Stopping growth is not a substitute but rather an addition to these methods to increase product titre, yield and productivity.

## Materials and methods

### Bacterial strains, plasmids, and growth medium

Strains and plasmids are listed in Table 1. *E. coli* DH5α was used for plasmid cloning and propagation. Genomic alterations and plasmid-based switcher experiments were performed in *E. coli* MG1655. *E. coli* was grown in lysogeny broth (LB) supplemented with the appropriate amount of antibiotics (100 μg/mL ampicillin, 25 μg/mL chloramphenicol, 25 μg/mL kanamycin, 10 μg/mL tetracycline) when necessary for the selection of strains and maintenance of the plasmids.

**Table 1.** Strains and plasmids.

| Strain | Description | Source/Reference |
| --- | --- | --- |
| DH5α | F<sup>-</sup> *endA1 glnV44 thi-1 recA1 relA1 gyrA96 deoR nupG purB20 ϕ80dlacZΔM15 $\Delta$(lacZYA-argF)U169, hsdR17(*rK<sup>-</sup>mK<sup>+</sup>*), λ<sup>-</sup>* | laboratory stock |
| MG1655 | K-12 F<sup>-</sup> λ<sup>-</sup> *ilvG<sup>-</sup> rfb-50 rph-1* | laboratory stock |
| bAJ78 | MG1655 ΔoriC::cat-attP-oriC-attB, Cm<sup>R</sup> | This study |
| bAJ83 | bAJ78/pAJ27, Cm<sup>R</sup> Kan<sup>R</sup> | This study |
| bAJ84 | bAJ78/pAJ35, Cm<sup>R</sup> Kan<sup>R</sup> | This study |
| bAJ85 | bAJ78/pAJ27/pAJ144, Cm<sup>R</sup> Kan<sup>R</sup> Amp<sup>R</sup> | This study |
| bAJ86 | bAJ78/pAJ35/pAJ144, Cm<sup>R</sup> Kan<sup>R</sup> Amp<sup>R</sup> | This study |
| **Plasmid** |  |  |
| pAJ27 | Kan<sup>R</sup>, P<sub>lacI</sub>-cl857, lambdaP<sub>R</sub>(T41C)-phiC31Int(1-29) in pBR322, negative control, truncated phiC31 integrase | This study |
| pAJ35 | Kan<sup>R</sup>, P<sub>lacI</sub>-cl857, lambdaP<sub>R</sub>(T41C)-phiC31Int in pBR322 | This study |
| pAJ144 | Tet<sup>R</sup>, LuxR-6His-mRFP1 in pJB866 | This study |
| pAJ157 | Amp<sup>R</sup>, P<sub>tac</sub>-Crimson in p15A | This study |
| pAJ181 | Kan<sup>R</sup>, P<sub>lacI</sub>-cl857, lambdaP<sub>R</sub>(T41C)-sulA in pBR322 | This study |
| pInt | Kan<sup>R</sup>, P<sub>lacZ</sub>-phiC31Int in pACYC177 | <sup>33</sup> |
| pKD46 | Amp<sup>R</sup>, lambda-Red helper plasmid (recombinase) | <sup>34</sup> |
| pKD3 | Amp<sup>R</sup> Cm<sup>R</sup>, template plasmid for FRT-flanked *cat* cassette | <sup>34</sup> |
| pTc-Wasabi | P<sub>lacI</sub>-cl857, P<sub>R</sub>/P<sub>L</sub>-mWasabi in pETDuet-1 | <sup>35</sup> |

Cm<sup>R</sup>, chloramphenicol resistance; Kan<sup>R</sup>, kanamycin resistance; Amp<sup>R</sup>, ampicillin resistance; Tet<sup>R</sup>, tetracycline resistance

### DNA manipulations

Short oligonucleotide sequences for cloning and sequencing were ordered from Metabion International AG.

*E. coli* MG1655 genomic *in situ* engineering was performed using recombineering. The Lambda Red recombination system was expressed from the pKD46 plasmid ^34^. Synthetic terminators (L3S3P25, L3S2P24 ^36^), *attB* and *attP* sites and tac promoter (without operators, Table 2) were ordered as synthetic DNA from Twist Bioscience. Flanking 400 bp long homologous regions were amplified from the genomic DNA of *E. coli* MG1655. The chloramphenicol resistance marker with adjacent FRT sites was amplified from pKD3. Recombineering fragment f_bAJ78 (Figure S1a, File S6) was assembled using overlap extension PCR, gel purified, and transformed into *E. coli* cells by electroporation.

**Table 2.**
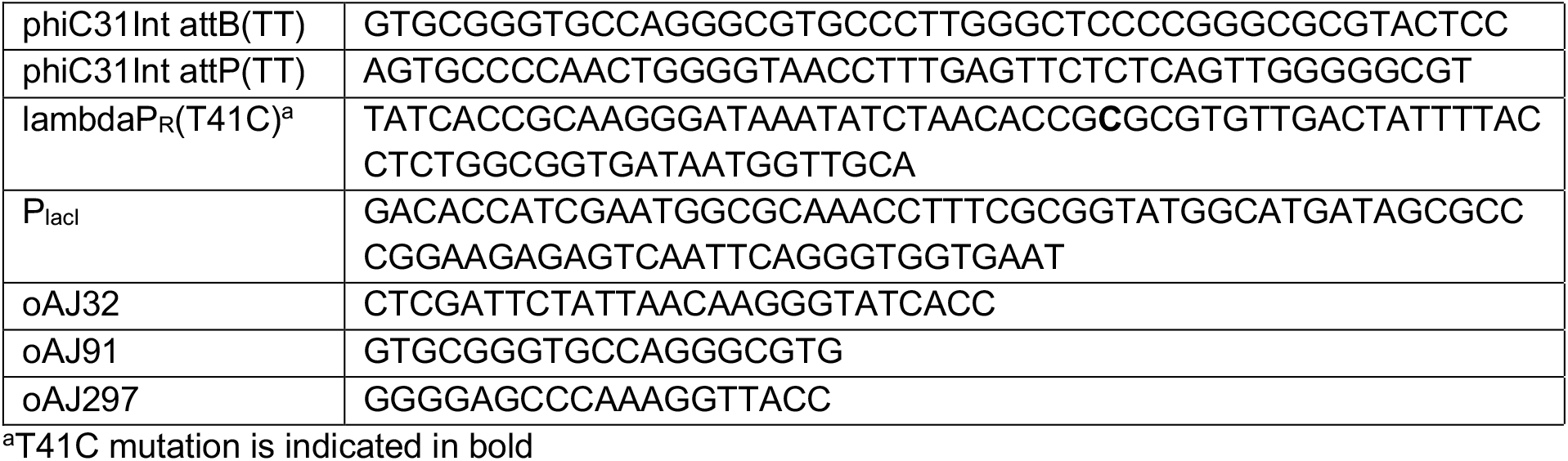
The nucleotide sequence of functional elements and switch-specific testing primers.

Plasmids were constructed using the CPEC method ^37^.

Plasmid pAJ35 (Figure S1b, File S1) was constructed to express phiC31 integrase and cI857 repressor in the switcher strain. In the plasmid, the DNA sequence of phiC31 integrase ^19^ was amplified from pInt plasmid and placed under the control of a mutated lambda P_R_(T41C) promoter (Table 2) ^18^. The P_R_(T41C) promoter is regulated by the cI857 repressor. A temperature-sensitive mutant of phage lambda repressor cI587 ^18^ together with the constitutive P_lacI_ promoter (Table 2) was amplified from pTcI-Wasabi plasmid. pInt plasmid was a gift from Michele Calos (Addgene plasmid # 18941 ; http://n2t.net/addgene:18941 ; RRID:Addgene_18941) and pTcI-Wasabi was a gift from Mikhail Shapiro (Addgene plasmid # 86101 ; http://n2t.net/addgene:86101 ; RRID:Addgene_86101).

Control strains carry plasmid pAJ27 (Figure S1b, File S2), which contains all the same elements as pAJ35, except that phiC31 integrase ORF was truncated to translate only the first 29 amino acids (MDTYAGAYDRQSRERENSSAASPATQRSA).

For the determination of protein synthesis activity, a gene encoding red fluorescent protein mRFP1 ^38^ was placed under the control of homoserine-lactone-inducible (HSL-inducible) Lux promoter in plasmid pAJ144 (Figure S1b, File S7). In plasmid pAJ157 (Figure S1b, File S8) the gene encoding the fluorescent protein Crimson was placed under the control of IPTG-inducible tac promoter.

For the expression of SulA protein, the phiC31-integrase encoding sequence in pAJ35 was replaced with the *sulA* gene from the *E. coli* MG1655 genome, forming plasmid pAJ181 (Figure S1b, File S9).

### Validation of oriC excision

Cultures of switcher and control strains were grown overnight at 30°C, diluted 2000 times into 20 mL of fresh LB and grown at 30°C or two hours. Thereafter, the temperature was shifted to 37°C. Samples were taken hourly to count colony-forming units per volume of culture (CFU/mL) on LB-agar plates, and to analyse the excision of *oriC* by PCR. Specific primer pairs were used to detect either excised (oAJ297 and oAJ32) or retained (unexcised) (oAJ91 and oAJ32) versions of *oriC*. The DNA sequence of excision site of *oriC* was verified by sequencing.

### Measurement of growth and fluorescence

Cultures of switcher and control strains were grown overnight at 30°C, diluted into LB as indicated in the figure legends, and grown on a 96-well plate at 30°C for a further 4 hours until the temperature was shifted to 37°C (timepoint 0 h). Where appropriate, mRFP1 or Crimson synthesis was induced by adding 12 mM HSL or 100 mM IPTG, respectively (final concentrations), either immediately prior to the temperature shift or at indicated timepoints. The growth curves and mRFP1- or Crimson-related fluorescence (reflecting protein level) were determined using a BioTek Synergy MX plate reader. The optical density at 600 nm and fluorescence intensity (excitation at 584/13.5 nm and emission at 607/13.5 nm to detect mRFP1, or excitation at 611/20 nm and emission at 650/20 nm for detecting Crimson) were measured in a culture volume of 100 μL.

### Fluorescence microscopy

Samples for microscopy were collected from 96-well plate reader experiment five hours after the switch. Of each culture, 0.7 mL was pipetted onto a piece of 1% LB-agarose pad. Imaging was performed with a Zeiss Observer Z1 microscope with a 100 × /1.4 oil immersion objective and Axiocam 506 mono camera (Zeiss). Phase-contrast and mRFP1-fluorescence (excitation 540–580 nm; emission 615–675 nm) images were recorded.

### Time-lapse microscopy

Cultures of switcher and control strains were grown overnight at 30°C, diluted 100 times into 3 mL of fresh LB and grown at 30°C for 2 hours. Of each culture, 0.7 mL was pipetted into a glass bottom dish (Thermo Scientific, 150682), covered with a piece of approximately 4 × 4 × 1 mm 1% LB-agarose pad. A cover glass was placed on top of each pad, and the dish was covered with a lid to minimize drying. The prepared cells were observed using a Zeiss Observer Z1 microscope with a 100 × /1.4 oil immersion objective and Axiocam 506 mono camera (Zeiss). The switch was induced under the microscope from the beginning of imaging by maintaining the temperature of the agarose pad at 37°C using a Tempcontrol 37–2 digital (PeCon). Images were taken every two minutes for 6 hours. Multiple positions were imaged in one experiment using an automated stage and ZEN software (Zeiss). The signals from GFPmut2 (excitation 460–490 nm; emission 509– 550 nm) and mRFP1 (excitation 540–580 nm; emission 615–675 nm), together with phase contrast, were recorded.

### Label-free proteomic analysis

#### 1) Growth conditions

Triplicate cultures of switcher and control strains were pre-grown overnight in sterile MOPS medium ^39^ with 0.3% glucose (MOPSglucose) at 30°C and diluted 800 times into fresh MOPSglucose at the start of the experiment. Cultures were grown for 8 hours at 30°C and thereafter 28 h at 37°C. Samples were collected 4–7 h after the temperature shift (growing phase, OD ≈ 1.0), 11–17 h after the temperature shift (early plateau, OD ≈ 2.3–3.0) and 28 h after the temperature shift (late plateau, OD ≈ 2.0–2.8). To collect samples from the switcher low plateau, triplicates of switcher strains were grown overnight in sterile MOPSglucose and diluted 2400 times into fresh MOPSglucose. Cultures were grown for 8 hours at 30°C, then 11.5 h at 37°C. Low-plateau samples were collected 11.5 h after the temperature shift. At indicated timepoints, 1 mg dry weight of cells calculated from OD at 600 nm ^40^ were harvested by centrifugation at 600 RCF for 7 minutes. The samples were then washed with PBS, pelleted by centrifugation again, and the pellets were flash-frozen in liquid nitrogen and stored at -80°C.

#### 2) Proteomics sample preparation and nano-LC-MS/MS analysis

Sample preparation was conducted using methods described ^41^. Frozen bacterial pellets were thawed and resuspended in lysis buffer (4% SDS, 100 mM Tris, pH 7.5, 10 mM dithiothreitol), heated at 95°C for 5 min, and sonicated. The protein concentration was determined by tryptophan fluorescence, and 30 μg of total protein was loaded into 30-kDa-cutoff Vivacon 500 ultrafiltration spin columns (Sartorius). Samples were digested for 4 h on a filter with 1:50 Lys-C (Wako) and thereafter overnight with 1:50 proteomics-grade dimethylated trypsin (Sigma-Aldrich) as described for the filter-aided sample preparation protocol ^42^. Peptides were desalted using C18 StageTips ^43^, eluted, dried, and reconstituted in 0.5% trifluoroacetic acid. Nano-liquid chromatography with tandem mass spectrometry (LC-MS/MS) analysis was performed as described previously ^44^ using an Ultimate 3000 RSLCnano system (Dionex) and a Q Exactive mass spectrometer (Thermo Fisher Scientific) operating with top-10 data-dependent acquisition.

#### 3) MS raw data processing

Mass spectrometric raw files were analysed using MaxQuant software ^45^ package v. 1.5.6.5 (File S10). Label-free quantification with the MaxQuant LFQ algorithm was enabled with default settings. Methionine oxidation, glutamine/asparagine deamidation, and protein N-terminal acetylation were set as variable modifications, while cysteine carbamidomethylation was defined as a fixed modification. The search was performed against the UniProt (www.uniprot.org) *Escherichia coli* K12 reference proteome database (September 2015 version) using the tryptic digestion rule (including cleavages after proline). Only identifications of at least seven-amino-acid-long peptides were accepted and transfer of identifications between runs was enabled. Protein quantification criteria were set to one peptide with a minimum of two MS1 scans per peptide. Peptide-spectrum match and protein false discovery rate (FDR) were kept below 1% using a target-decoy approach. All other parameters were default. Summed peptide peak areas (protein intensities) were normalized using the MaxLFQ algorithm ^46^.

#### 4) Proteomics data analysis

Proteomics data were analysed using Perseus software ^47^ v. 1.6.15.0. Normalized LFQ intensity values were used as the quantitative measure of protein abundance. Protein identifications classified as “Only identified by site” and “Contaminants” were excluded from further analysis. LFQ intensity values of the whole proteome of each sample were used to conduct principal component analysis.

Protein LFQ intensity values were log2 transformed ^48^ and normal data distribution was verified from histogram distribution plots of log2 transformed data for each sample (data not shown). Samples were allocated into groups (two strains, four conditions: control growing, control early plateau, control late plateau, switcher growing, switcher early plateau, switcher late plateau, switcher low plateau).

For hierarchical clustering analysis, only proteins with complete data for all 21 samples were included. First, proteins with statistically significant changes in abundance between sample groups were identified using a multiple-sample ANOVA test, with p-values adjusted for multiple testing by the Benjamini–Hochberg permutation FDR at 1%. Statistically significant proteins were subjected to Z-score normalization. Hierarchical clustering analysis using Euclidean distances was performed and presented as a heatmap using Perseus software.

Proteins with three valid values in at least one sample group were used for further analysis. A two-way Student’s t-test was used to compare sample groups (Benjamini– Hochberg FDR < 0.01) (File S10). Of the statistically significant proteins, proteins with more than a twofold difference between LFQ intensities (|log2(LFQ a/LFQ b)| > 1) were interpreted as biologically significant.

The list of genes for differentially expressed proteins was used for enrichment analysis. Enrichment analysis for pathways was conducted using the SmartTables function of Pathway Tools v. 19.0 (available at biocyc.org) ^49,50^ with a threshold of Fisher’s exact test (p < 0.05).

## Acknowledgements

This work was supported by the European Union from the European Regional Development Fund through the Centre of Excellence in Molecular Cell Engineering (2014-2020.4.01.15-0013).

## Author contribution

A. Jõers conceived the study. All authors designed and performed experiments and analysed the data. A. Jõers, M. Kasari and V. Kasari wrote the manuscript. All authors read and approved the final version.

## Data availability

The mass spectrometry proteomics data have been deposited to the ProteomeXchange Consortium via the PRIDE ^51^ partner repository with the dataset identifier PXD029931 and 10.6019/PXD029931.

