## Supporting information for "Decoupling growth and production by removing the origin of replication from a bacterial chromosome"

The SI includes Figures S1-S5

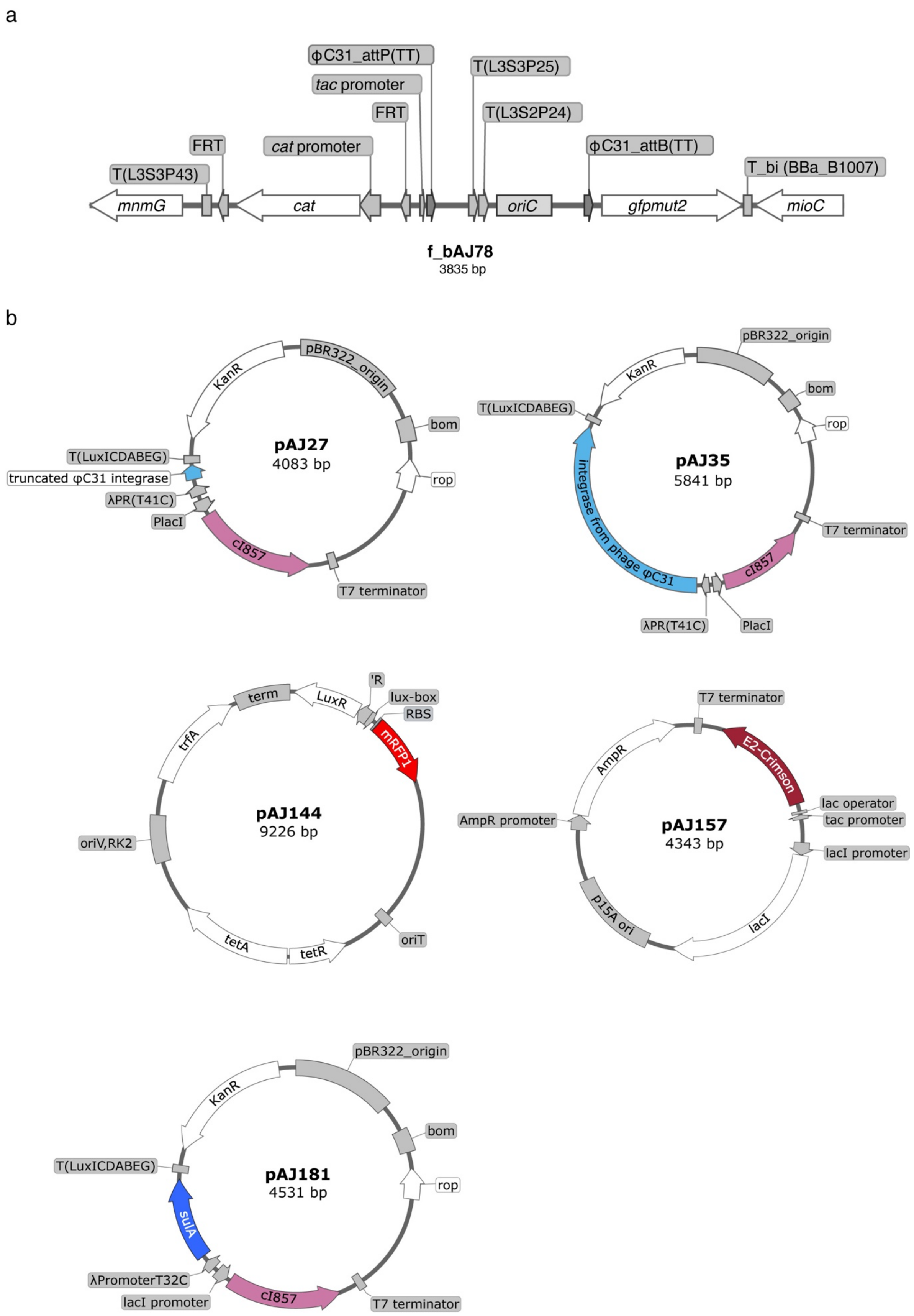


**Figure S1**. **Schematic maps of switcher constructs.** (**a**) Modified *oriC* genomic region in the genome of *E. coli* MG1655. (**b**) Maps of plasmids constructed in this study.

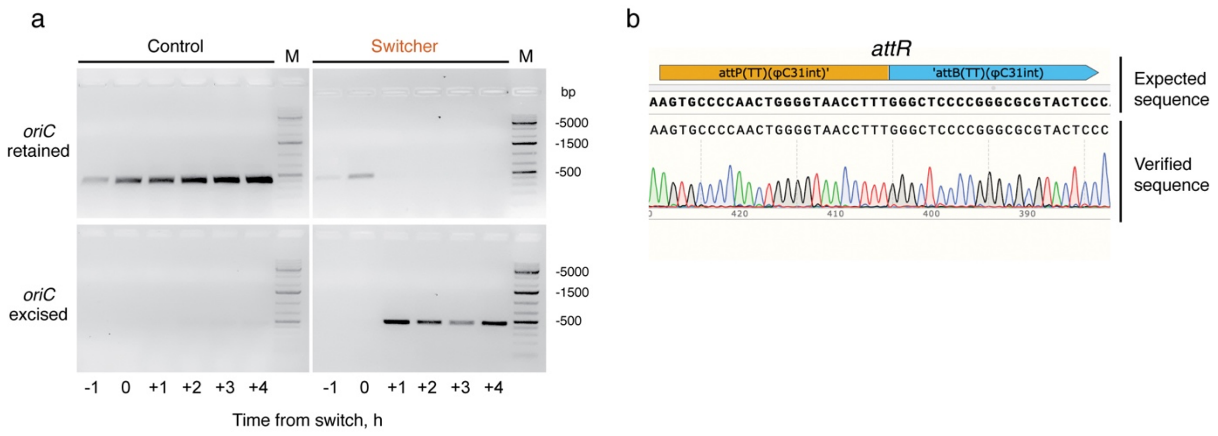


**Figure S2**. **Sequence verification of the switched cells.** (**a**) Uncropped gel images of PCR testing presented on Figure 1c. (**b**) Sequence chromogram of the *attR* junction formed after the switch.

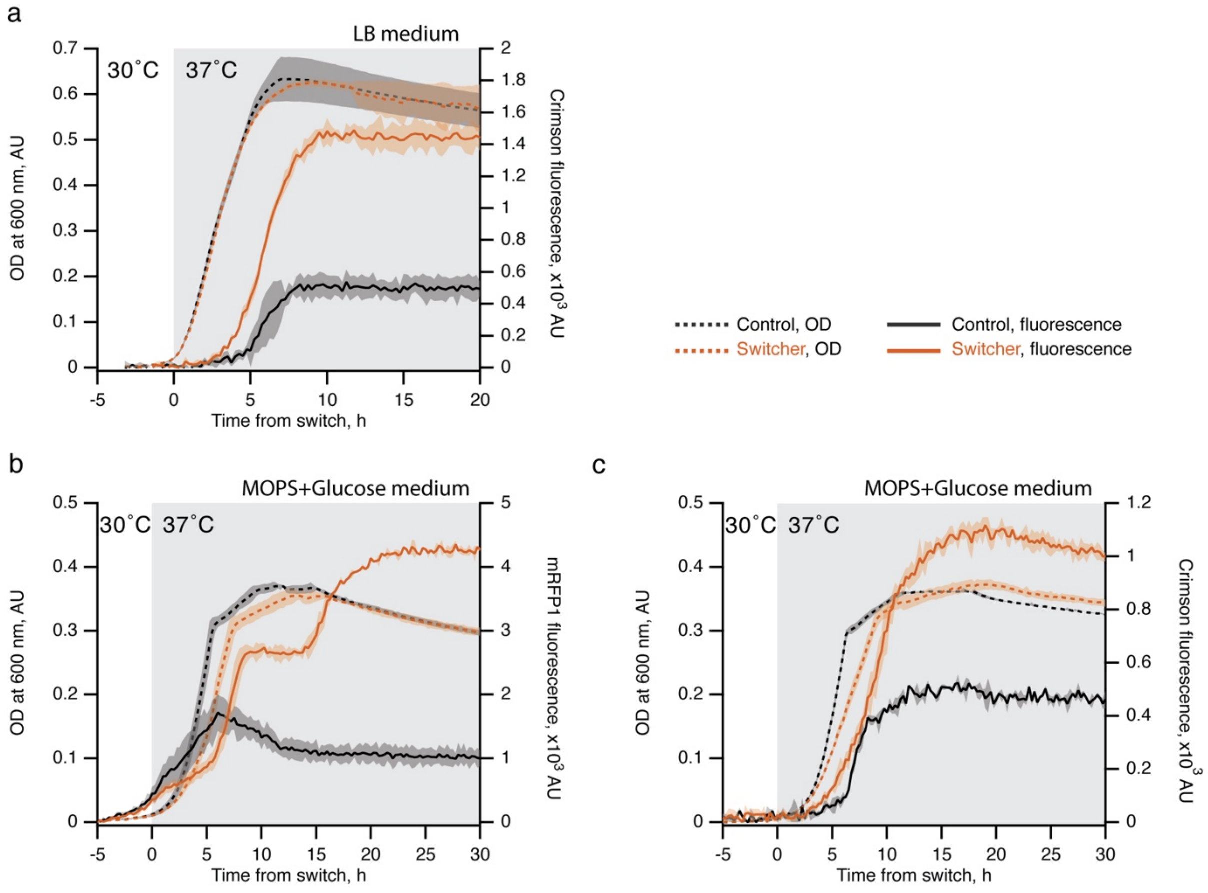


**Figure S3**. **Protein synthesis capacity of the switcher tested in different settings.** Control (black) and switcher (orange) cultures were pre-grown in LB (**a**) or MOPSglucose medium (**b**, **c**) at 30°C on a 96-well plate. At timepoint zero hours the temperature was changed to 37°C to induce the switching. The production of Crimson (**a**, **c**) and mRFP1 (**b**) proteins was induced by adding IPTG or homoserine lactone (HSL) at timepoint zero, respectively. In each experiment, the optical density at 600 nm was monitored. The mean of three biological replicates is plotted, shading indicates standard deviation.

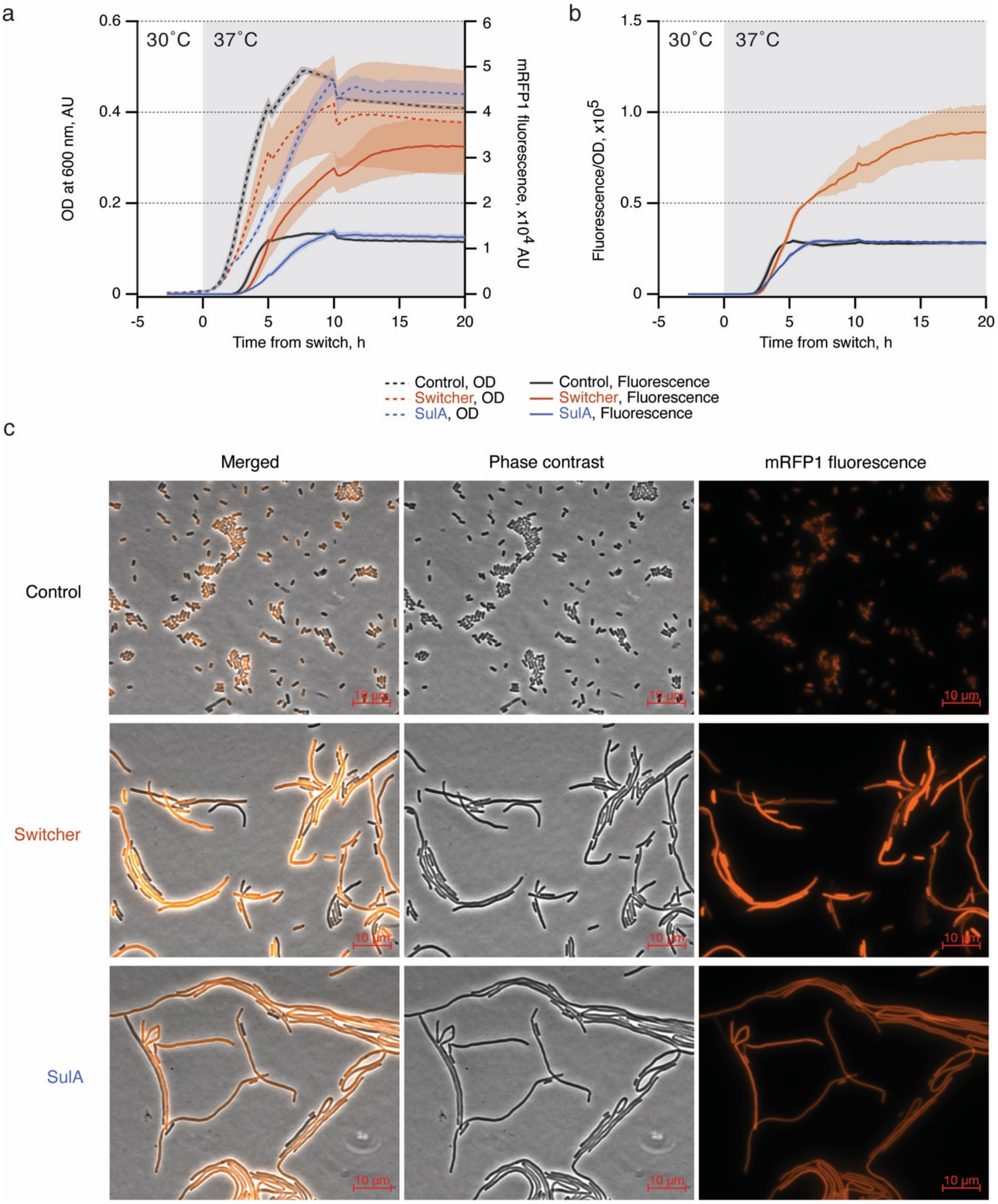


**Figure S4. SulA-mediated cell elongation does not increase protein production efficiency.** (**a**) Control (black), switcher (orange), and SulA (blue) cultures were pre-grown in LB at 30°C on a 96-well plate. At timepoint zero hours the temperature was changed to 37°C to induce the switching or the expression of *sulA*. Production of mRFP1 protein was induced by adding homoserine lactone (HSL) at timepoint zero. The optical density at 600 nm and fluorescence at 607 nm were monitored. (**b**) The fluorescence-over-optical-density ratio was calculated based on the values in panel **a** and plotted. The mean of three biological replicates is plotted. Shading indicates standard deviation. (**c**) Samples for microscopy were collected from the plate reader experiment five hours after the switch in panel **a**. Phase contrast and mRFP1-fluorescence images were captured and representative images are presented here.

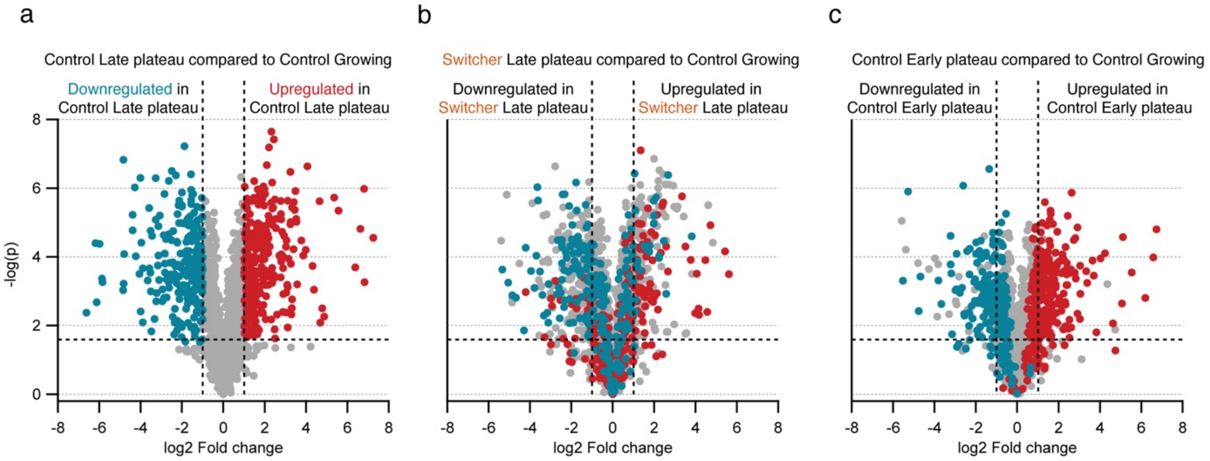


**Figure S5**. **Distribution of up- and downregulated proteins in late-plateau cells.** (**a**) Comparison of protein expression levels between control growing and control late-plateau samples. Red dots represent proteins upregulated in late-plateau cells, blue dots represent the downregulated ones (fold change > 2, false discovery rate < 1%). (**b**) Comparison of protein expression levels between control growing and switcher late-plateau samples. Protein colour coding is the same as in panel **a**. (**c**) Comparison of protein expression levels between control growing and control early-plateau samples. Protein colour coding is the same as in panel **a**.
